# Adhesion of dry and wet cribellate capture silk

**DOI:** 10.1101/160408

**Authors:** Hervé Elettro, Sébastien Neukirch, Arnaud Antkowiak, Fritz Vollrath

## Abstract

We demonstrate the impressive adhesive qualities of Uloborid spider orbweb capture when dry, which are lost when the nano-filament threads are wetted. A force sensor with a 50 nN–1mN detection sensitively allowed us to measure quantitatively the stress–strain characteristics of native silk threads in both the original dry state and after wetting by controlled application of water mist with droplet sizes ranging between 3 and 5 μm and densities ranging between 10^4^ and 10^5^ per mm^3^. Stress forces of between 1 and 5 μN/μm^2^ in the native, dry multifilament thread puffs were reduced to between 0.1 and 0.5 μN/μm^2^ in the wetted collapsed state, with strain displacements reducing from between 2 and 5 mm in the dry to 0.10–0.12mm in the wetted states. We conclude that wetting cribellate threads reduce their van der Waals adhesion with implications on the thread’s adhesive strength under tension. This should be considered when discussing the evolutionary transitions of capture silks from the ancestral dry-state nanofilaments of the cribellate spider taxa to the wet-state glue droplets of the ecribellate taxa.

## Introduction

Orb-web spiders use two very different mechanisms to entrap insects in their capture threads. The evolutionary more ancestral cribellate technique requires the spider to slowly and laboriously hackle thousands of fine filaments while the more advanced (i.e. derived) ecribellate technology deploys highly cost-efficient self assembling glue droplets [Vollrath 2005]. Molecular profiling suggests that the two mechanisms have dissimilar evolutionary histories with very strong evidence for ‘independent origins for the two types of orb webs’ [Bond et al 2014; Fernández et al 2014] despite many similarities in architecture and ecology [Shear 1986; Bond and Opell 1998; Foelix 2011; Opell and Bond 2011]. The two opposing mechanisms of prey capture rely on fundamental differences between the two types of silk used by the two weaver types, cribellate and ecribellate, in their orb-web capture threads.

To briefly recap: Orb web-building spiders are using two functionally opposing prey capture systems [Peters 1987; Vollrath 2005; Opell and Schwend 2009, Sani et al 2011] i.e. the hackled-and-puffed nano-thread-adhesion capture system of the cribellate spiders [Peters 1984; Opell et al. 1994] and the two-component-extrusion glue-adhesion system of the ecribellate spiders [Vollrath and Edmonds 1989; Opell and Hendricks 2007]. The hackled threads require the spider to comb and electrostatically charge the threads, which are understood to attach and adhere to the insect by van-der-Waals forces [Hawthorn and Opell 2002, 2003]. The aqueous glue threads carry droplets that self-assemble via a Rayleigh-Plateau transition upon water adsorption from the atmosphere [Edmonds and Vollrath 1992], and attach by surface wetting and glycoprotein adhesion [Vollrath et al 1990, Vollrath and Tillinghast 1991; Opell and Hendricks 2007]. Consequently it has been deduced that the glue threads only work when wet while hackled threads work best when dry [Peters 1987; Opell 1994] or ‘dryish’ [Hawthorn and Opell 2003].

It seems from the literature [cited so far as well as see also e.g. Blackledge and Hayashi 2006] that the cribellate system would fundamentally depend on its original, non-wetted, highly ‘puffed out’ configuration state for the nano-fibrils to retain their function. Indeed, the very spinning mechanism of the cribellum fibre composite is specially adapted to an electrostatic spinning process that leads to the configuration of hackled puffs of dry silk nano-filaments astride core carrier threads [Kronenberger and Vollrath 2015]. One must argue that the puffs, in turn, would rely on dryness for their continued function, if indeed electrostatic forces and nano-adhesion sites are key to their functionality. Confusingly, is has also been shown that high ambient humidity seems to increase the adhesive properties of some (nano-noded) cribellum threads, perhaps by adding capillary forces to electrostatic forces [Hawthorn and Opell 2003], strange as that might sound considering potentially conflicting physical dynamics.

Here we test the hypothesis that cribellate capture threads are indeed much more ‘sticky’ when dry as opposed to when wetted. Uloborus cribellum threads were exposed to high-density mist and their adhesion to a nano-force tensile tester measured before and after wetting. Imagery of both dry/puffed and wetted/collapsed threads complemented the force measurements.

## Results and Discussion

Our experiment demonstrates that wetting destroys the adhesive properties of the threads. We conclude that ‘watering’ renders the *Uloborus* capture threads unable to retain any prey that they may intercept (Figure 1).

**FIGURE 1.**
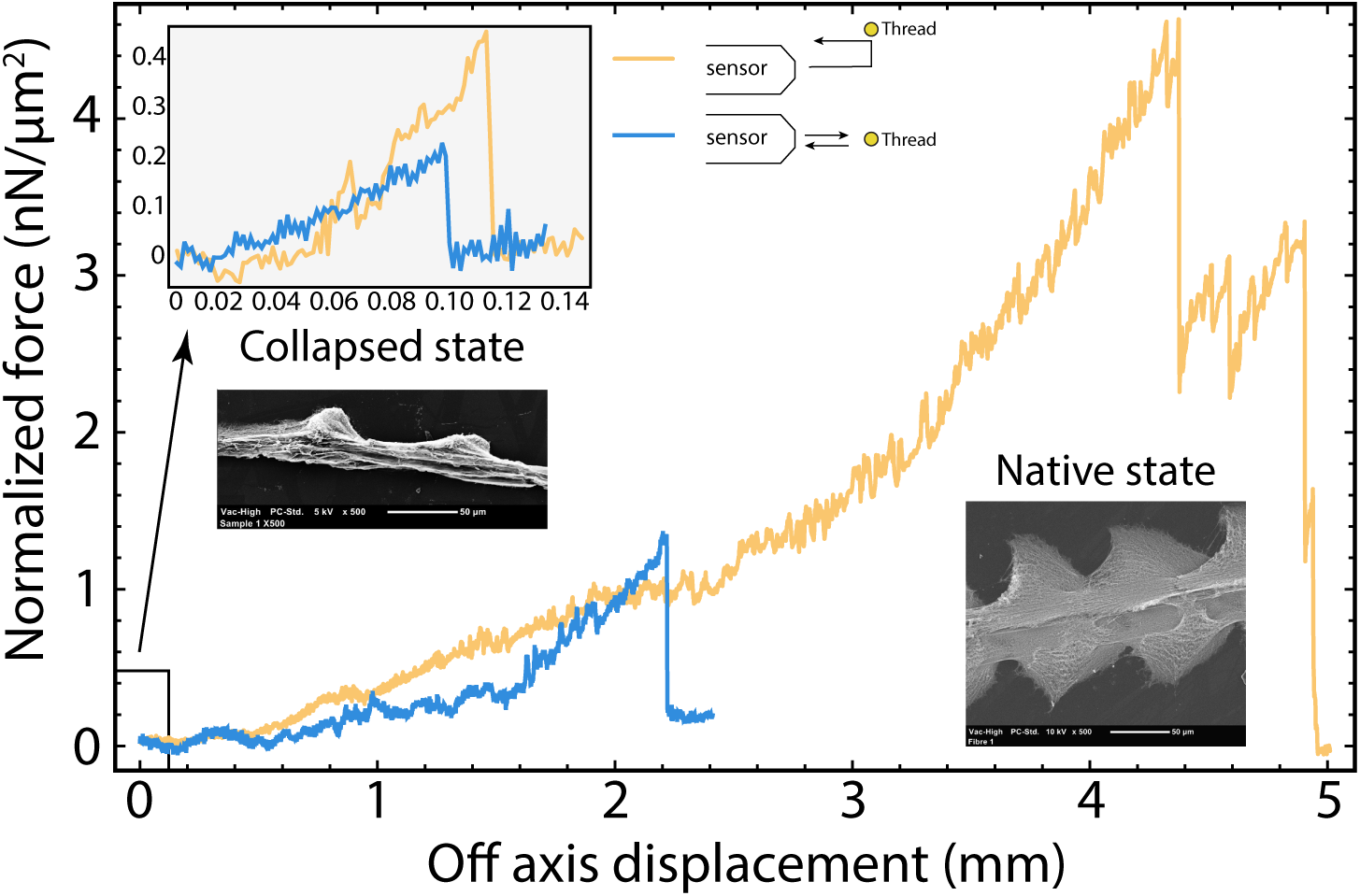
Representative stress-stain graphs of native and, collapsed cribellate capture silk adhering to a sensor. Measurements were taken subsequently on the same piece of capture thread of *Uloborus plumipe* and images were taken from adjacent sections of the same thread. The sensor was either pushed into the thread or lowered onto the thread before being pulled away, in both cases the force of contact was equivalent. Please note that these two curves are representative for 14 individual stress-strain tests, further explanations and significances in the text. Inserts: SEM images of a capture thread in the native puffed state and collapsed after wetting.

*Uloborus plumipes* spiders were collected at the Paris Jardin des Plantes greenhouse and taken into the laboratory where they spun webs in appropriate frames. The experimental apparatus consisted of a combination of microscope and stress-strain gage with the added ability of controlled application of water mist. *Uloborus* capture thread samples were carefully taken from a web using calipers to avoid deformation, straining and stressing. The samples were then tested using a FemtoTools, FT-FS1000 detection capacitive force sensor with a 50nN-1mN measurement range in two different ways (press-in and press-on) reflecting the ways an insect might approach the web(forward impact and lateral ‘flapping’). Wetting was achieved by controlled mega-sonic misting of distilled water at controlled droplet sizes ranging between 3-5 μm and at densities ranging between 10^4^ and 10^5^ per mm^3^. We note (i) that threads dried out very quickly (see also video in Suppl. Material and/or www-link) and (ii) that it made no difference to dimensions and adhesive properties whether prior to the measurements pre-wetted threads were left at room humidity (50% rH) or thoroughly dried over P2O5. SEM images were taken at a range of magnifications with 10nm of Gold/Palladium coating.

Our observations demonstrated that, in its native state, the cribellate silk studied showed the typical uloborid puffs. These collapsed during even brief 5min wetting (see Suppl Materials and www-link for video). The measured adhesion forces differed significantly between native-puffed and wetted-collapsed threads (Fig 1) and this was irrespective whether the sensor was pushed forward into the capture thread, or pushed down unto it (one tailed t-tests, p = 1.0% and 0.75% resp., N=4, n=14). For the dry threads pushing onto a thread followed by pulling away showed stronger adhesion than pushing into a thread, again this was highly significant (Fig. 1, one tailed t-test p= 1.4%). We assign this difference in force to the concurrent difference in contact between the sensor and the individual filaments. In the case of push-in/pull-out the contact area of the sensor would have been about 2500μm^2^ and adherent threads were pulled away at more or less 90 degree. In the case of push-down/pull-away the contact area of the sensor would have been about 5000μm^2^ and the threads are pulled in the area of contact and in a very oblique angle, which would allowed for much longer periods of contact over the same pulling distance. Of course, the differences of actual filament contact area between sensor and threads would change when the filaments are all collapsed into one-another, as happens when they are wetted (Fig 1 left inset), which is in stark contrast to their native state when they are fully puffed out (Fig1 right inset).

The experimental wetting may or may not have affected electrostatic charges of the thread by temporary ‘grounding’ of the otherwise insolating threads via the applied aqueous mist coating. However, it is much more likely that the wetting-induced collapse of the filament puffs significantly decreased the number of surface contact area/points and that this alone would account for the significant drop in adhesion after wetting. As our images show, as Peters [1987] predicted and as Zheng et al [2010] confirmed, wetting the submicron capture filaments of *Uloborus* causes them to coalesce, which is an important phenomenon in fibre physics and hence reasonably well understood [Bico et al 2004]. As we have shown here, fine mist accumulates to quickly and effectively destroy the adhesive effectiveness of hackled silk. For the spider this means that fog (or perhaps already dew) may radically decrease the capture efficiency of an *Uloborus* orb web. As is well known, all orb weavers, cribellate and ecribellate alike, tend to take down their webs in rain. In the ecribellates this is a response to droplet overloading, which leads to sagging and snapping threads that can compromise the integrity of the whole structure. As we have now demonstrated, in the ecribellates this is likely to be in response to loss of function.

## Acknowledgements

We all thank the Royal Society of London (International Exchange grant IE130506), the Paris group thanks the Agence Nationale Reseaux (grant 09-JCJC-0022-01), La Ville de Paris (grant Programme Emergence) and the CNRS (grant PEPS PTI) while FV thanks the US Air Force (AFOSR grant FA9550-12-1-0294) and the European Research Council (ERC grant SP2-GA-2008-233409). **Data accessibility:** All data and methods are reported within this paper and with the electronic supplementary material.

## SUPPLEMENTARY VIDEO

Mist is sent onto cribellate capture thread, revealing the uloborid puffs in the first instants, before collapsing them into non-sticky spindle-knots. This reduced significantly the adhesion properties of the capture thread, effectively destroying its primary biological function.

## References

Bico J, Roman B. Moulin L, Boudaoud A (2004) Elastocapillary coalescence in wet hair. Nature 432, 690.

Blackledge TA, Hayashi CY (2006) Unraveling the mechanical properties of composite silk threads spun by cribellate orb-weaving spiders. J. Exp. Bio. 209, 3131–3140.

Bond JA, Opell DB (1998) Testing adaptive radiation and key innovation hypotheses in spiders. Evolution 52, 403–414.

Bond JE, Garrison NL, Hamilton CA, Godwin, RL, Hedin M, Agnarsson I (2014) Phylogenomics Resolves a Spider Backbone Phylogeny and Rejects a Prevailing Paradigm for Orb Web Evolution. Curr. Biol. 24, 1765–1771.

Edmonds D, Vollrath F (1992) The contribution of atmospheric water vapour to the formation and efficiency of a spider’s capture web. Proc. Roy. Soc. London 248, 145–148.

Fernández R, Hormiga G, Giribet G (2014) Phylogenomic Analysis of Spiders Reveals Nonmonophyly of Orb Weavers. Curr. Biol. 24, 1772–1777.

Foelix R (2011) Biology of Spiders. 3rd ed, Oxford, Oxford Univ. Press.

Hawthorn AC, Opell BD (2002) Evolution of adhesive mechanisms, in cribellar spider prey capture thread: evidence for van der Waals and hygroscopic forces. Biol. J. Linn. Soc. Lond. 77, 1–8.

Hawthorn, AC, Opell BD (2003) Van der Waals and hygroscopic forces of adhesion generated by spider capture threads. J. Exp. Biol. 206, 3905–3911.

Kronenberger K, Vollrath F (2015) Spiders spinning electrically charged nano-fibres. Biol. Let. 11(1):20140813. doi: 10.1098/rsbl.2014.0813.

Opell BD, Tran AM, Karinshak SE (2011) Adhesive compatibility of cribellar and viscous prey capture threads and its implication for the evolution of orb-weaving, spiders. J. Exp. Zool. 315, 376–384.

Opell BD (1994) Factors governing the stickiness of cribellar prey capture threads in the spider family Uloboridae. J. Morphol. 221, 111–119.

Opell DB, Hendricks ML (2007) Adhesive recruitment by the viscous capture threads of, araneoid orb-weaving spiders. J. Exp. Biol. 210, 553–560.

Opell BD, Schwend HS (2007) The effect of insect surface features on the adhesion of, viscous capture threads spun by orb-weaving spiders. J. Exp. Biol. 210, 2352–2360.

Opell BD, Schwend HS (2008) Adhesive efficiency of spider prey capture threads. Zoology 112: 16–26.

Opell DB, Bond JE (2001) Changes in the mechanical properties of capture threads and the evolution of modern orb-weaving spiders. Evol. Ecol. Res. 3, 567–581.

Peters TM 1984 The spinning apparatus of Uloboridae in relation, to the structure and construction of capture threads, (Arachnida, Araneida). Zoomorph. 104:96–104

Peters HM 1987 Fine Structure and Function of Capture Threads in Ecophysiology of Spiders. pp 187-202 in (ed.) W. Nentwig Springer-Verlag Berlin Heidelberg

Peters, H. M. (1995). Ultrastructure of orb spiders’ gluey capture threads. Naturwissenschaften 82, 380–382.

Sahni V, Blackledge TA, Dhinojwala A (2011) A review on spider silk adhesion. J. Adhes. 87, 595–614.

Shear W (1986) Spiders, Webs, Behavior and Evolution. Stanford University Press

Vollrath F (2005) Spiders’ Webs. Curr. Biol. 15, R364–365.

Vollrath F, Fairbrother WJ, Williams RJP, Tillinghast EK, Bernstein DT, Gallager KS, Townley MA (1990) Compounds in the droplets of the orb spider’s viscid spiral. Nature 345, 526–528.

Vollrath F, Edmonds D (1989) Modulation of the mechanical properties of spider silk by coating with water. Nature 340, 305–307.

Vollrath F, Tillinghast E. (1991) Glycoprotein glue beneath a spider web’s aqueous coat. Naturwissenschaften 78, 557–559.

Zheng Y, Bai H, Huang Z, Tian X, Nie FQ, Zhao Y, Zhai J, Jiang L (2010) Directional water collection on wetted spider silk. Nature 462, 640–443.

